# Reversing chemorefraction in colorectal cancer cells by controlling mucin secretion

**DOI:** 10.1101/2021.09.17.460827

**Authors:** G Cantero-Recasens, J Alonso-Marañón, T Lobo-Jarne, M Garrido, M Iglesias, L Espinosa, V Malhotra

## Abstract

15% of colorectal cancers (CRC) cells exhibit a mucin hypersecretory phenotype, which is suggested to provide resistance to immune surveillance and chemotherapy. We now formally show that colorectal cancer cells build a barrier to chemotherapeutics by increasing mucins’ secretion. We show that low levels of KChIP3, a negative regulator of mucin secretion (Cantero-Recasens *et al*., 2018), is a risk factor for CRC patients’ relapse in subset of untreated tumours. Our results also reveal that cells depleted of KChIP3 are four times more resistant (measured as cell viability and DNA damage) to chemotherapeutics 5-Fluorouracil plus Irinotecan (5-FU+iri.) compared to control cells, whereas KChIP3 overexpressing cells are 10 times more sensitive to killing by chemotherapeutics. Similar increase in tumour cell death is observed upon chemical inhibition of mucin secretion by the sodium/calcium exchanger (NCX) blockers (Mitrovic *et al*., 2013). Finally, sensitivity of CRC patient-derived organoids to 5-FU+iri increases 40-fold upon mucin secretion inhibition. Reducing mucin secretion thus provides a means to control chemoresistance of mucinous colorectal cancer cells and other mucinous tumours.

## INTRODUCTION

Secreted mucins are synthesized in the Endoplasmic Reticulum (ER) and transported to the Golgi apparatus, where they receive extensive O-glycosylation that increases their molecular weight by about 5-fold. These heavily glycosylated mucins are then packed into specialized micrometre-sized granules where they can reach molecular weights of up to 50 MDa (1, 2). These granules mature by a process that involves mucin condensation and, finally, a small number of granules fuse to plasma membrane in a calcium dependent manner to release the required quantity of mucins in the extracellular space (3, 4). Specifically, mucins are secreted by two main processes: 1) extracellular calcium dependent pathway called stimulated secretion, which is triggered by an exogenous agonist; and 2) intracellular calcium dependent release, but independent of external agonists, called baseline secretion. It has recently been shown that cooperation between TRPM4/5 sodium channels and the Na^+^/Ca^2+^ exchangers (NCX) control stimulated secretion (5, 6). The calcium binding protein KChIP3, on the other hand, regulates baseline secretion (7).

The secreted mucins compose the mucus that protects the underlying epithelium from toxins, pathogens and allergens (8). Secretion of the right quantity and quality of mucins is crucial because mucin hypersecretion in the airways is linked to chronic obstructive diseases and hyposecretion makes the underlying tissue more susceptible to damage by the immune cells and the foreign particles (2, 9). Most cancer cells start expressing mucins and it is widely thought that they isolate tumour cells from the immune system and also render them resistant to anti-cancer therapies (10). For instance, patients with mucinous colorectal adenocarcinoma (10-15% of CRC (11)) present dysregulated mucin secretion that is long-associated with high resistance to chemotherapy. In fact, extracellular mucins account for at least 50% of the volume of these tumours (12). Whether or not controlling the levels of mucins secreted by cancer cells affects their susceptibility to chemotherapeutics, however, remains untested. Here we have studied this question directly in CRC cell lines and organoids derived from CRC patients and show that reducing mucin secretion reverses the chemorefractory properties of hypersecretory CRC cells.

## RESULTS

### 5-FU+iri. (5-Fluorouracil+Irinotecan) treatment induces mucin production

15-20% of colorectal cancers (CRC) develop a mucinous profile (11), which renders cancer cells highly resistant to commonly used chemotherapies, such as 5-Fluorouracil plus Irinotecan (5-FU+iri.) (13). To study the contribution of secreted mucins to CRC chemorefraction, we used the HT29 CRC cell line (clones HT29-18N2 and HT29-M6) that are differentiated into mucin producing cells (5, 14).

We first tested the effect of 5-FU+iri. on mucin production, focusing on secreted mucins, which are the main macrocomponents of the mucus layer (i.e. MUC5AC) (2). Briefly, differentiated HT29-M6 cells were treated with 5-FU+iri. (50 μg/mL 5-Fluorouracil + 20 μg/mL Irinotecan) and MUC5AC levels monitored over time (0, 1, 3, 6 and 24 hours) by western blot (WB). Our results reveal that 5-FU+iri. treatment strongly induces MUC5AC production (**Figure 1A**). To further confirm this effect, 5-FU+iri. treated HT29-M6 cells (24 hours) were imaged by confocal immunofluorescence microscopy. Accordingly, we found that 5-FU+iri. treated cells present a clear increase in MUC5AC level compared to control cells (**Figure 1B**). These results suggest that increased secretion of mucins after 5-FU+iri. treatment could act as a barrier to chemical permeation thus enhancing chemorefraction of mucosecretory cancer subtypes.

**Figure 1.**
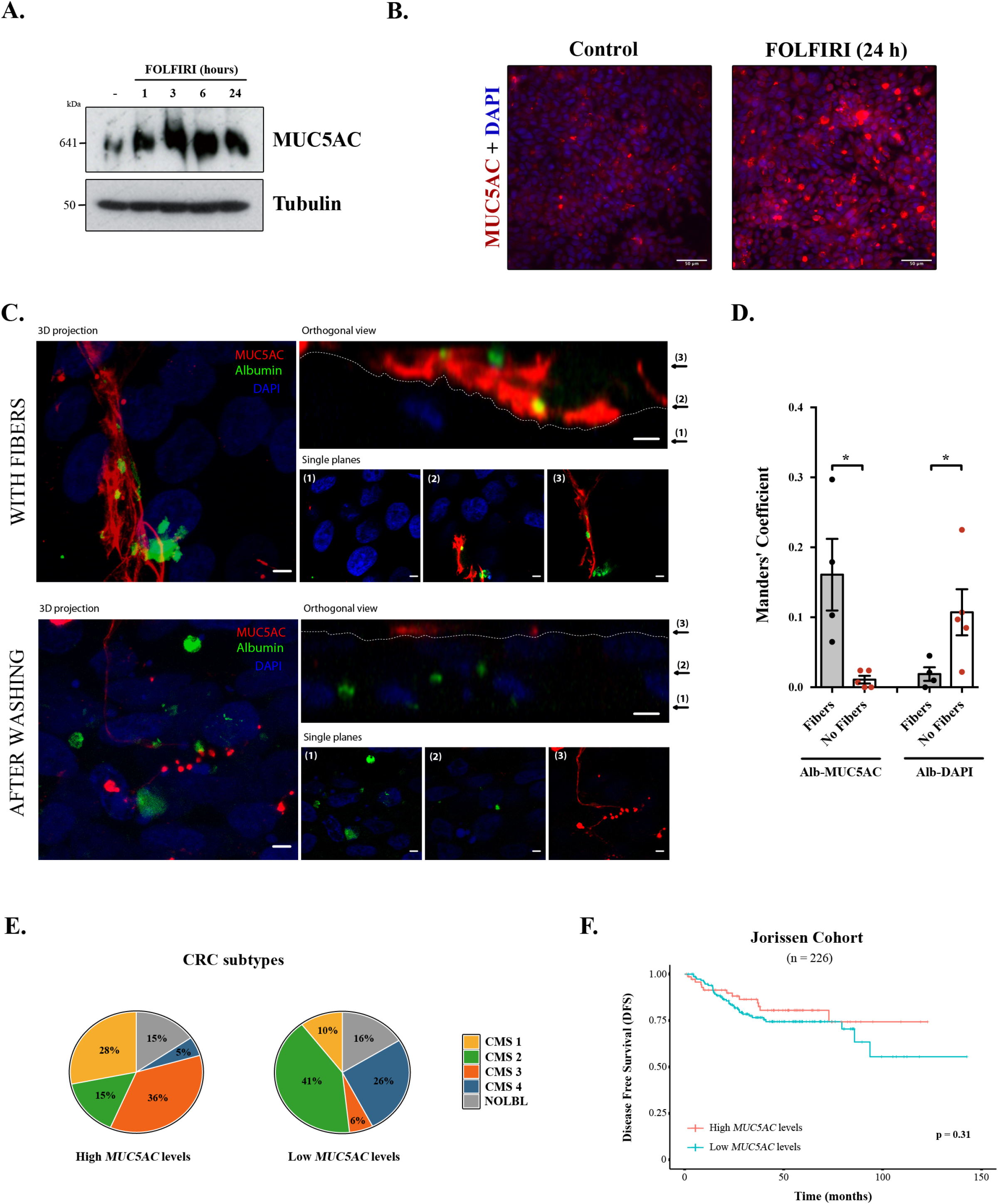
Mucins in colorectal cancer. **A)** Cell lysates from differentiated HT29-M6 cells treated with 5-FU+iri. (50 µg/mL 5-FU + 20 µg/mL iri.) for 24h were analysed by western blot with an anti-MUC5AC to test expression levels. Tubulin was used as a loading control. **B)** Immunofluorescence Z-stack projections of differentiated HT29-M6 cells treated with vehicle (Control) or 5-FU+iri. (50 µg/mL 5-FU + 20 µg/mL Irinotecan) for 24h. Cells were stained with anti-MUC5AC (red, and DAPI (blue). Scale bar = 50 µm. **C)** Differentiated HT29 cells were treated with 100 µM ATP for 30 minutes, washed extensively (upper panel) or smoothly (lower panel), and incubated with Alexa Fluor 488-labelled albumin for one hour. Cells were stained with anti-MUC5AC antibody (Red) and DAPI (Blue). In the orthogonal view, dotted lines across the images demarcate the top surface of the cell. Scale bars correspond to 5 µm. **D)** Colocalization between MUC5AC and Albumin or DAPI and albumin was calculated from immunofluorescence images by Manders’ coefficient using FIJI. Average values ± SEM are plotted as scatter plot with bar graph. The y-axis represents Manders’ coefficient of the fraction of albumin overlapping with MUC5AC or DAPI. **E)** Consensus molecular subtype (CMS) distribution of high and low MUC5AC expressing colorectal tumours (Jorissen cohort). **F)** Disease free survival (DFS) of colorectal cancer patients with high (n=73) or low (n=153) MUC5AC levels (Jorissen cohort). Abbreviations: CMS: Consensus Molecular Subtype, NOLBL: Samples without a defined CMS subtype (tumours with No Label). *p<0.05, **p<0.01.

Differentiated HT29-M6 cells were seeded in glass bottom dishes and treated with 100 µM ATP, a well-known secretagogue for mucins (15). After 30 minutes, cells were divided in two groups: one group was extensively washed with isotonic solution to remove the mucin fibres, while the other group was subjected to mild washes to preserve these fibres. Next, cells were incubated with Alexa Fluor 488-labelled albumin for one hour, processed for confocal immunofluorescence microscopy and stained with anti-MUC5AC antibody. As shown in figure 1C, mucins act as a barrier that prevents the endocytosis of albumin by HT29-M6 cells (**Figure 1C**, upper panel). Contrarily, removal of mucin fibres allows the contact and uptake of albumin by cells (**Figure 1C**, lower panel). Quantification of the images using the FIJI software revealed that in the presence of mucin fibres, albumin colocalizes with MUC5AC (Manders’ coefficient Albumin-MUC5AC: 0.16 vs. 0.01, p=0.01); whereas in cells without mucin barrier, albumin is found intracellularly (Manders’ coefficient Albumin-DAPI: 0.02 vs. 0.11, p=0.03) (**Figure 1D**). Altogether, these data demonstrate that colonic cancer cells produce higher levels of MUC5AC in response to chemotherapy treatment, which form a barrier that protects cells from incorporation of extracellular compounds. We anticipated that MUC5AC may act as a barrier against CT in human CRC tumours and as a consequence it could represent a prognostic biomarker for CRC.

Analysis of CRC subtypes, using the consensus molecular subgroup -CMS-classification from Guinney and collaborators (16) indicated a radically different distribution of high and low MUC5AC expressing tumours among subtypes: CMS1 (28% vs. 10%), CMS2 (15% vs. 41%), CMS3 (36% vs. 6%) and CMS4 (5% vs. 26%) (**Figure 1E**). Importantly, both CMS1 (associated with immune evasion) and CMS3 (metabolic dysregulation and goblet cell markers enrichment) have worse prognosis than CMS2 (16), which include the majority of tumours with low MUC5AC expression. These results point to an important role of MUC5AC in the prognosis of CRC patients. To directly assess this possibility, we evaluated whether MUC5AC expression levels discriminates patients’ outcome in a well-annotated CRC cohort (GSE14333, Jorissen cohort (17)). Disease free survival (DFS) analysis revealed no significant differences between patients with high or low MUC5AC levels (**Figure 1F**) strongly suggesting that factors controlling mucin secretion, and not only mucin levels, might be relevant biomarkers for CRC prognosis.

### The mucin regulator KChIP3 is a risk factor for CRC with high MUC5AC expression

Mucin secretion can be triggered by an external agonist (<u>stimulated secretion</u>), such as ATP and allergens; or, in the absence of external agonists (<u>baseline secretion</u>) (3). We have shown previously that KChIP3, an intracellular calcium sensor, negatively controls baseline mucin secretion (loss of KChIP3 causes mucin hyper-secretion) (7). So we checked whether KChIP3 could be a prognostic biomarker for CRC. Analysis of DSF of patients according to KChIP3 levels, encoded by the *KCNIP3* gene, revealed that KChIP3 is, indeed, a prognostic factor (HR=1.925, p=0.02) (**Figure 2A**). To further confirm whether this effect is related to mucins, patients were divided between high (n=73) and low (n=153) MUC5AC expression levels, and then we studied the effect of KChIP3 on the disease-free survival (DFS) of patients. Our analysis reveals that low levels of KChIP3 are a clear risk factor in those patients with high MUC5AC levels (HR=3.29, p=0.04) (**Figure 2B**), whereas there is no significant effect on patients with low MUC5AC expression (**Figure 2C**). These results suggest that low levels of KChIP3, which consequently increase the quantity of mucins secreted, protect cancer cells from chemotherapeutic drugs.

**Figure 2.**
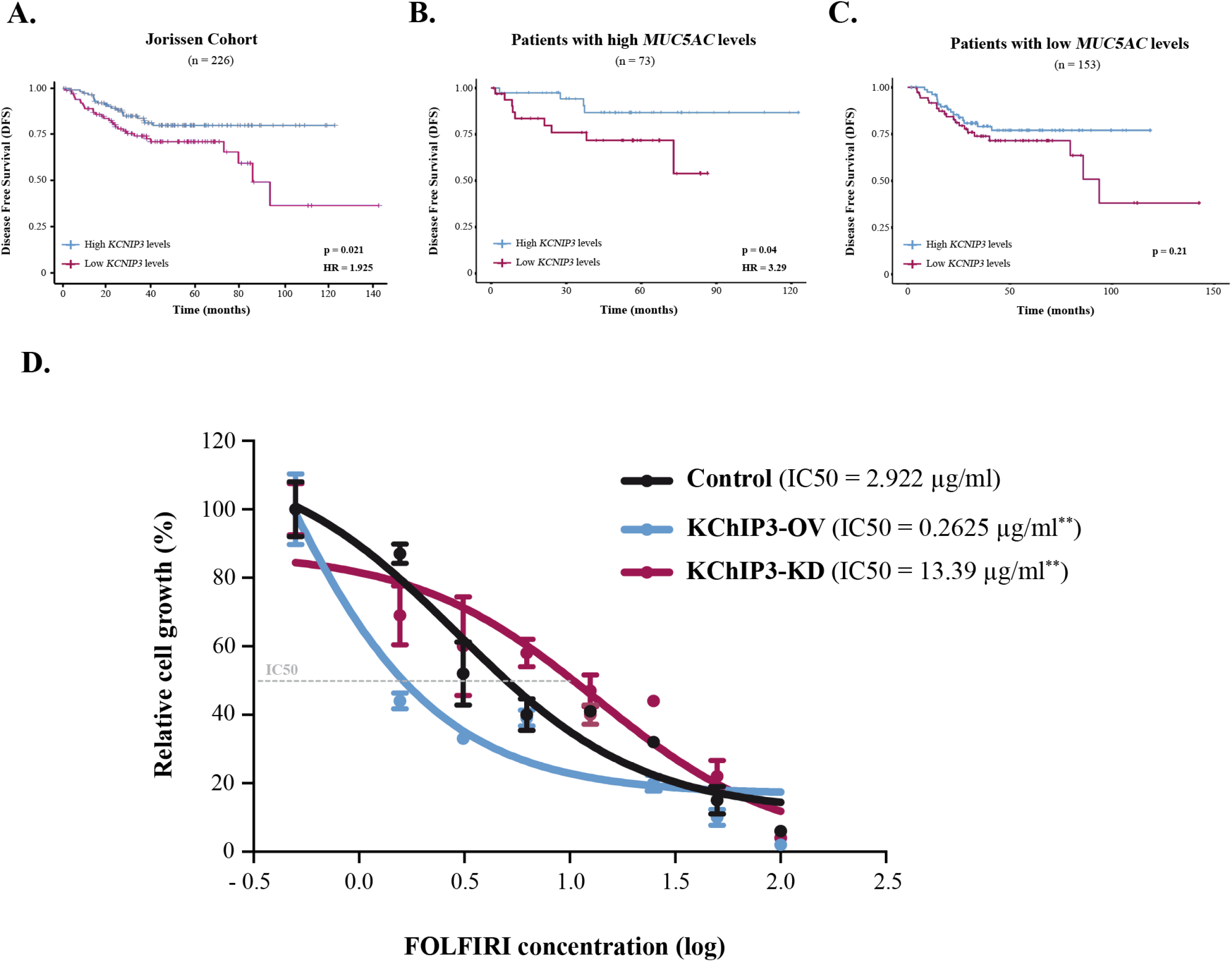
KChIP3 is a prognostic marker for CRC. **A-C)** Disease free survival (DFS) according to KChIP3 levels of colorectal cancer patients (low KChIP3 levels, n=120; high KChIP3 levels, n=106) **(A)**, CRC patients with high MUC5AC levels (low KChIP3 levels, n=39; high KChIP3 levels, n=34) **(B)** or CRC with low MUC5AC levels (low KChIP3 levels, n=81; high KChIP3 levels, n=72) **(C). D)** Differentiated control (black), KChIP3 overexpressing cells (KCNIP3-OV, blue) and KChIP3 depleted cells (KCNIP3-KD, red) were treated for 72 hours with increasing concentrations of 5-FU+iri. Average values ± SEM are plotted as scatter plot. The y-axis represents the percentage of cell growth relative to the lowest concentration of 5-FU+iri. The IC50 was calculated from the interpolated curve. Abbreviations: HR: Hazard Risk, 5-FU+iri.: 5-FU+iri. *p<0.05, **p<0.01.

### Modulation of KChIP3 levels alters CRC cells chemoresistance

To further test this hypothesis and the role of KChIP3 in chemoresistance, we used HT29-18N2 cell lines stably depleted or overexpressing KChIP3 (KChIP3-KD or KChIP3-OV, respectively) (7). Briefly, differentiated control, KChIP3-KD (increased mucin secretion) and KChIP3-OV (reduced mucin release) cells were treated with increasing concentrations of 5-FU+iri. and then monitored for cell viability using the CellTiter-Glo assay. As shown in figure 2D, cells depleted of KChIP3 are 4-fold more resistant to 5-FU+iri. than control cells, whereas overexpression of KChIP3 renders cancer cells almost 10 times more sensitive to the chemotherapy (IC50 control cells= 2.9 µg/ml, KChIP3-KD cells= 13.4 µg/ml, KChIP3-OV= 0.3 µg/ml) (**Figure 2D**).

In summary, low levels of KChIP3 increase mucin secretion (7), which protects CRC cells from chemotherapy. Contrarily, high levels of KChIP3 inhibit baseline mucin release thereby reducing the mucus barrier and rendering cancer cells more sensitive to 5-FU+iri.

### Inhibition of MUC5AC secretion enhances the effect of chemotherapy

Although baseline mucin secretion and its regulator KChIP3 have a clear role in the response to chemotherapy of CRC cells, our results indicated that 5-FU+iri. treatment can trigger mucin production. Interestingly, it was previously described that stimulated mucin secretion depends on extracellular calcium entry through TRPM4/5 – Na^+^/Ca^2+^ exchangers (NCX) cooperation (6). We anticipated that if CT-induced mucin secretion was increasing CRC chemoresistance, Benzamil, an NCX inhibitor (18), should enhance the effect of 5-FU+iri. on HT29-M6 CRC cells. Differentiated HT29-M6 cells were treated with 5-FU+iri. (IC50) and 20 µM Benzamil or vehicle, and the levels of DNA damage were determined by the standard comet assay (19). Our results showed that co-treatment with 5-FU+iri. and Benzamil increased the levels of DNA breaks compared to 5-FU+iri. plus vehicle (**Figure 3A**). We then tested cell viability of HT29-M6 at 24 and 72 hours post-treatment of 5-FU+iri. alone or in combination with Benzamil. As shown in figure 3B, there was no significant effect of the combination treatment at 24 hours. Importantly, the therapeutic effect of low concentration of 5-FU+iri., which reduced cell viability around 15% at 72 hours, was strongly enhanced when combined with 20 µM Benzamil (40% reduction on cell viability compared with the control) (**Figure 3B**). These results demonstrate that treatment with the NCX inhibitor Benzamil improves the effect of chemotherapy on tumour cell eradication.

**Figure 3.**
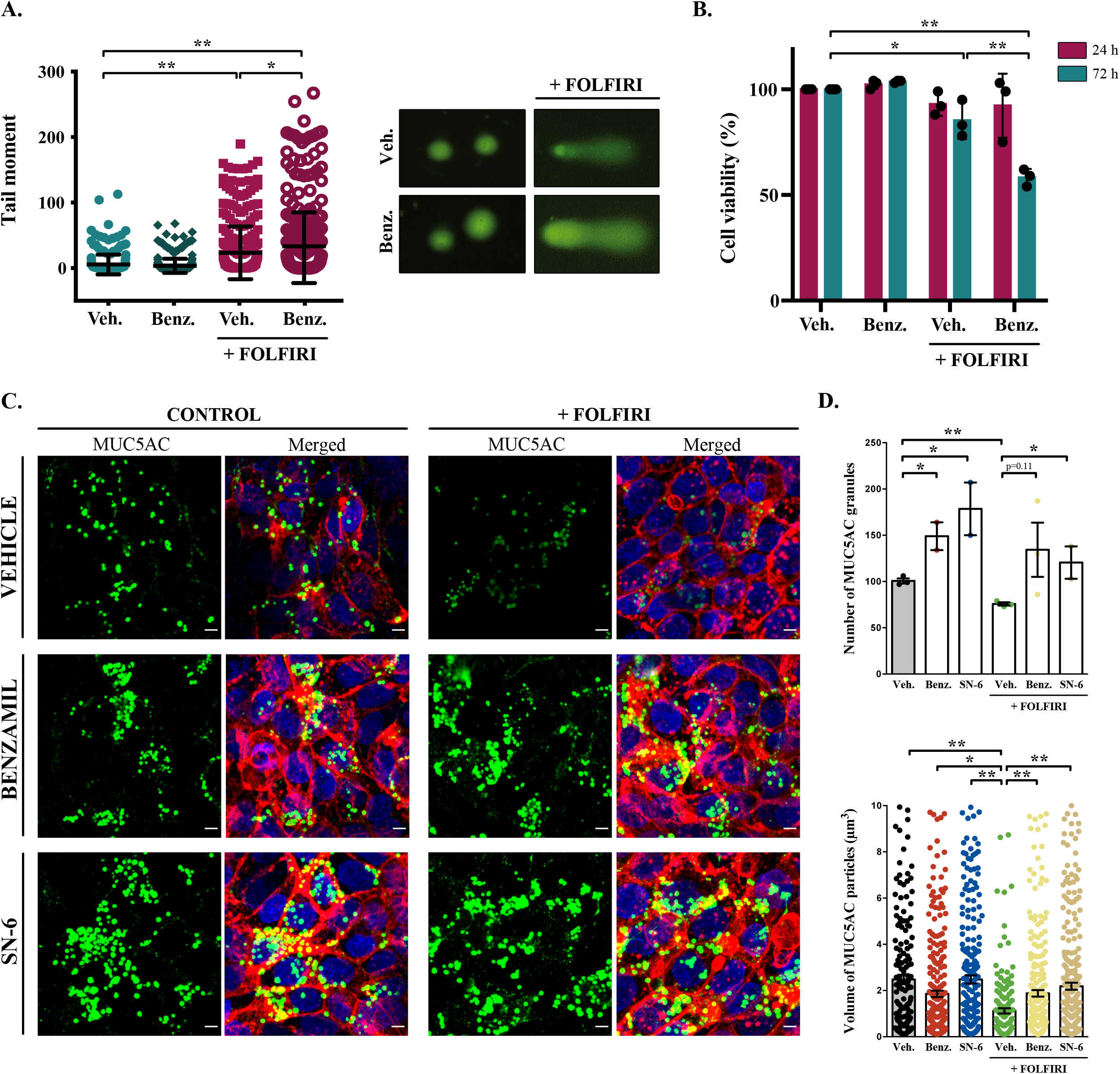
Inhibition of NCXs enhances cell death by 5-FU+iri. **A)** Comet assay of HT29-M6 cell line treated with 5-FU+iri. (25 µg/mL 5-FU + 10 µg/mL Irinotecan) and 20 µM Benzamil, alone or in combination. The tail moment was measured after 72 hours of treatment. **B)** Quantification of cell viability in HT29-M6 cell line after treatment (24 or 72 hours) with 5-FU+iri. (10 µg/mL 5-FU + 4 µg/mL Irinotecan) and 20 µM Benzamil, alone or in combination. C) Immunofluorescence Z-stack projections of differentiated HT29-M6 cells treated with vehicle, 20 µM Benzamil or 10 µM SN-6 in presence or absence of 5-FU+iri. Cells were stained with anti-MUC5AC (green), phalloidin (red) and DAPI (blue). Scale bar = 5 µm. D) Quantification of the number (upper graph) and volume (lower graph) of MUC5AC granules from immunofluorescence images by confocal microscope. Average values ± SEM are plotted as scatter plot with bar graph. Abbreviations: Veh.: Vehicle, Benz.: Benzamil. *p<0.05, **p<0.01.

To further confirm that this additive effect was due to the inhibition of 5-FU+iri.-dependent mucin secretion, we used a second inhibitor called SN-6, which is a specific blocker of the reverse mode (calcium influx) of NCX and can be used at lower concentration, therefore presenting fewer side effects than Benzamil (20). HT29-M6 cells were grown at post-confluence (differentiation) in glass-bottom dishes and then left untreated or treated with 5-FU+iri. (IC50) in combination with vehicle, 20 µM Benzamil or 10 µM SN-6. Next, cells were processed for immunofluorescence with anti-MUC5AC antibody and imaged by confocal microscopy to visualize mucin granules (**Figure 3C**). Analyses of images using FIJI software (21) revealed that cells treated with 5-FU+iri. contained significantly fewer MUC5AC-containing particles than control cells (75.7 *vs*. 100.7 particles/field, p=0.002) with a reduced average size (1.12 µm^3^ vs. 2.48 µm^3^) (**Figure 3C**, quantification in **Figure 3D**), which was in agreement with enhanced mucin secretion as shown in figure 1. Importantly, co-treatment with Benzamil or SN-6 reverted this effect causing a strong accumulation of mucin granules, as shown by an increase in the number of MUC5AC-containing particles (Benzamil: 1.88 particles/field, SN-6: 2.19 particles/field) and their size (Benzamil: 134.3 µm^3^, SN-6: 120.5 µm^3^), compared to cells treated with 5-FU+iri. and vehicle (**Figure 3C**, quantification in **Figure 3D**). Interestingly, SN-6 had a stronger effect than Benzamil on cells treated with 5-FU+iri. further indicating that NCX inhibition enhanced sensitivity to chemotherapy.

### Specific inhibition of NCX enhances 5-FU+IRIdependent damage in both cells and patient-derived organoids

In order to test whether SN-6 effect observed on mucins correlated with increased sensitivity to 5-FU+iri. treatment, we determined the levels of γH2AX, a standard marker of genotoxic damage (22) in HT29-M6 cells treated with 5-FU+iri. (IC50) in presence or absence of 10 µM SN-6. We obtained cell lysates at different time points (0, 1, 3 and 6 hours) after treatment and then measured the levels of MUC5AC and γH2AX by western blot. Our results confirmed that SN-6 leads to MUC5AC intracellular accumulation over time in 5-FU+iri. treated cells (**Figure 4A, first lane**), which correlated with a slight increase in γH2AX levels (**Figure 4A, second lane**). To validate this increase in DNA damage, differentiated HT29-M6 cells were treated with vehicle (Control), 5-FU+iri. (IC50), 10 µM SN-6 or 5-FU+iri.+SN-6, and processed for immunofluorescence with anti-MUC5AC and anti-γH2AX antibodies (**Figure 4B**). Quantification of γH2AX positive cells confirmed the increase on DNA damage in cells treated with 5-FU+iri. and SN-6 (Control: 0.9 %, SN-6: 0.4 %, 5-FU+iri.: 8.0 %, 5-FU+iri. + SN-6: 24.8%) (**Figure 4C**).

**Figure 4.**
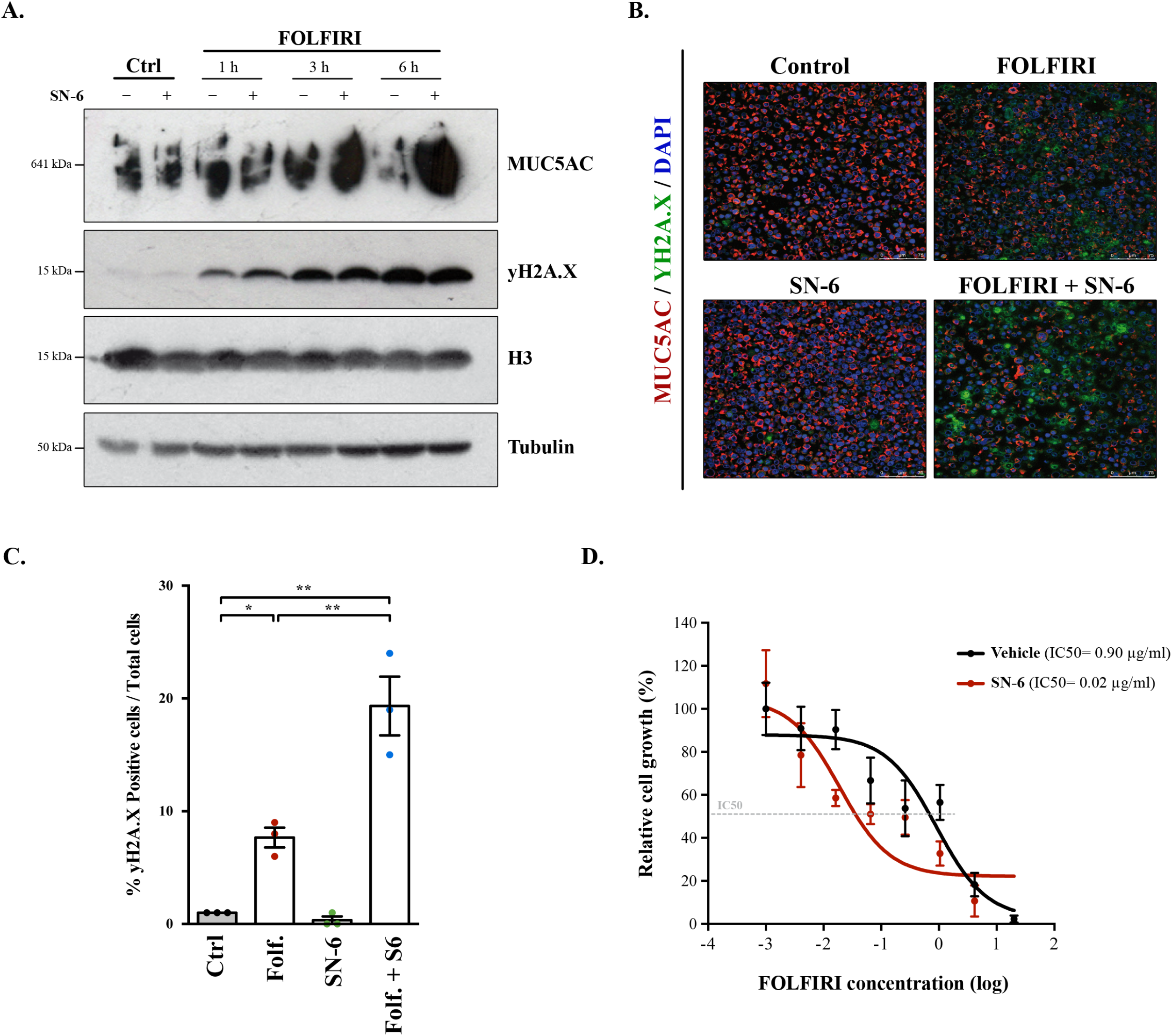
SN-6 treatment increases sensitivity of CRC derived cells and organoids to 5-FU+iri. **A)** Cell lysates of differentiated HT29-M6 cells pre-treated with 10 µM SN-6 inhibitor for 24h and then exposed to vehicle or 5-FU+iri. (50 µg/mL 5-FU + 20 µg/mL iri.) for 0, 1, 3 and 6 hours were analysed by western blot with an anti-MUC5AC, anti-yH2A.X and H3 to test their levels. Tubulin was used as a loading control. **B)** Immunofluorescence images of differentiated HT29-M6 cells treated with vehicle (control) or 10 µM SN-6 in presence or absence of 5-FU+iri. (50 µg/mL 5-FU + 20 µg/mL Irinotecan) for 24h. Cells were stained with anti-MUC5AC (red), anti-yH2A.X (green) and DAPI (blue). Scale bar = 50 µm. **C)** Quantification of the number of yH2A.X positive cells in the different conditions relative to the total number of cells from the immunofluorescence images. **D)** CRC patients derived organoids (PDOs) were treated for 72 hours with increasing concentrations of 5-FU+iri. with vehicle or 10 µM SN-6. Average values ± SEM are plotted as scatter plot. The y-axis represents the percentage of cell growth relative to the lowest concentration of 5-FU+iri. The IC50 was calculated from the interpolated curve. Abbreviations: Ctrl: control, Folf.: 5-FU+iri., Folf.+S6: 5-FU+iri. + SN-6. *p<0.05, **p<0.01.

Finally, to test the pathophysiologic relevance of our findings we studied the effect of SN-6 + 5-FU+iri. combination on a CRC patient-derived organoids (PDO). With this aim, we treated PDO cells, which we recently validated as model for therapy prescription (19), with suboptimal doses of 5-FU+Iri. alone or in combination with SN-6. Consistent with our cellular studies, mucins secretion inhibition with SN-6 increased up to 40 times the sensitivity of PDO cells to 5-FU+iri. compared with cells treated with vehicle (IC50 5-FU+iri. + vehicle = 0.90 µg/ml, IC50 5-FU+iri. + SN-6 = 0.02 µg/ml) (**Figure 4D**).

Together these results show that mucin secretion inhibitors alleviate the resistance to chemotherapeutics agents of cells that secrete mucins.

## DISCUSSION

Mucins are high molecular weight O-glycosylated proteins produced by goblet cells that line the epithelium of several organs of the airway and the digestive tract (3, 9). These proteins are the major macro-components of the mucus layer, which is a physical barrier that protects the underlying tissue from external insults (23). Importantly, the combination of different mucins and the type of secretion (baseline or stimulated) modulates the properties of the mucus layer to meet the tissue demands. For instance, baseline secretion is the prominent pathway that maintains the colonic mucus layer, while stimulated is the main pathway to rapidly protect the epithelium after allergen or toxic challenge in the airway (24). Accordingly, tight regulation of mucin quantities and composition is essential for the correct functioning of the mucus. Alterations in their synthesis or release are, therefore, hallmarks of several pathologic situations like severe asthma or colorectal cancer (CRC). Indeed, hyper production of mucin in some subtypes of CRC leads to enhanced chemoresistance of cancer cells thereby reducing patients’ survival (25, 26). But what is the contribution of mucins to CRC chemo-refraction? The chemicals and proteins employed here affect both MUC2 and MUC5AC as shown previously (6, 7), but for the sake of simplicity we have monitored only the effects on MUC5AC secretion.

### Cancer cells produce mucins as a protective response to chemotherapy

Our data show that CRC cells produce mucins in response to 5-Fluorouracil + Irinotecan (5-FU+iri.), a first-choice chemotherapy for most CRC (27). This is extremely important, because secreted mucins can create a barrier that could prevent drugs (for instance, chemotherapy) from reaching the tumour cells. Thus, cancer cells respond to treatment by secreting copious amounts of mucin to form a barrier to chemotherapeutics. An interesting question is how 5-FU+iri. triggers mucin secretion and whether there is any specific receptor that could be used as pharmacological targets.

### KChIP3, novel risk factor for mucinous CRC

We postulated that mucin levels could be used as prognostic biomarkers for CRC. Our analyses of a public CRC cohort, however, revealed that mucin levels do not reflect the prognosis of CRC patients. Although surprising, this is not totally unexpected because mucin secretion is an extremely well-regulated process to prevent pathological situations. Therefore, high intracellular levels do not directly translate to increased extracellular levels. Our data reveal that low levels of KChIP3, which negatively regulates baseline secretion (7), is an important risk factor for untreated tumours with high MUC5AC expression levels. We have found that cells with low levels of KChIP3 (that corresponds to more mucin secretion) are more resistant to 5-FU+iri. To summarise, low levels of KChIP3 promote mucin secretion, which is even more dramatic in tumours with high expression of MUC5AC. Increased mucin secretion protects cancer cells from chemotherapy, which drastically reduce CRC patients’ disease-free survival (DFS). KChIP3 thus emerges as a good prognostic marker, although it may have low relevance as a therapeutic target because it necessitates the use of gene therapy to enhance its expression and block mucin secretion. An exciting option is to target factors that control KChIP3 activity, such as ryanodine receptors or quenchers of intracellular calcium (7).

### Combined treatment with NCX inhibitors improves chemotherapy efficacy

Mucinous cancers present higher chemoresistance than other subtypes of CRC (11). As suggested by several authors and now shown by our experimental data, secreted mucins can act as a barrier to external insults including chemotherapeutic drugs. In fact, genetic regulation of this secretion by modulating KChIP3 levels renders cancer cells more sensitive to treatment. However, our results also demonstrate that 5-FU+iri. is a stimulus for mucin secretion, which may be an adaptive response of cancer cells for isolation from harmful environment. This unexpected effect can explain how some cancer cells become more resistant to treatment over time.

Stimulated mucin secretion depends on extracellular calcium entry through the reverse mode of sodium – calcium exchangers (NCX) (5, 6). Our data show that inhibition of stimulated mucin secretion reduces the resistance of tumour cells to anti-cancer treatments. Both NCX inhibitors (Benzamil and SN-6) prevent the effect of 5-FU+iri. on mucin secretion. On the other hand, co-treatment with NCX inhibitors improves the apoptotic effect of 5-FU+iri. on cancer cells, even at low 5-FU+iri. concentrations (IC20). However, one of the main problems of NCX inhibitors is their low specificity and the effects on cardiac function. To avoid these problems, SN-6 was developed as a specific inhibitor of NCX reverse mode that at working concentration does not affect other receptors, transporters, or ion channels; and has less side effects than Benzamil (20, 28). Our results reveal the potential therapeutic importance of SN-6. Finally, it is satisfying to note that the effect of SN-6 on the chemotherapy sensitivity of cancer cells is replicated in patients’ derived organoids.

## CONCLUSIONS

In conclusion, colorectal cancer cells produce mucins to create a protective environment against chemotherapy treatments. Blocking mucin secretion by genetic (altering KChIP3 levels) or chemical means (using NCX inhibitors) renders tumour cells more sensitive to anti-cancer therapies. In addition, we have identified the mucin secretion regulator KChIP3 as a novel risk factor for mucinous CRC, which could be used as a prognostic biomarker. We have provided potential new pharmacological strategies to control chemorefraction of mucinous colorectal cancers, but this approach is likely to benefit patients with mucinous cancers in general.

## MATERIALS AND METHODS

### Bioinformatics analysis

Transcriptomic and available clinical data dataset from CRC was downloaded from the open-access resource CANCERTOOL. We used the Jorissen cohort (GSE14333) which includes expression and clinical data for 226 patients (17). Patients were classified according to the mean expression of either MUC5AC or KChIP3. The association with relapse was assessed using Kaplan-Meier estimates and Cox proportional hazard models. A standard log-rank test was applied to assess significance between groups. This test was selected because it assumes the randomness of the possible censorship. All the survival analyses and graphs were performed with R using the survival (v.3.2-3) and survimer (v.0.4.8) packages and a p-value<0.05 was considered statistically significant.

### Reagents and Antibodies

All chemicals were obtained from Sigma-Aldrich (St. Louis, MO) except anti-MUC5AC antibody clone 45M1 (Neomarkers, Waltham, MA). Secondary antibodies for immunofluorescence microscopy were from Life Technologies.

### Cell lines

#### HT29-M6

HT29-M6 cells (obtained from ATCC) were grown in Dulbecco’s modified Eagle’s medium (Invitrogen) plus 10% foetal bovine serum (Biological Industries) and were maintained in 5% CO2 incubator at 37ºC. All experiments were performed with cells at postconfluency when they presented higher levels of mucins.

#### HT29-18N2

HT29-18N2 cells (obtained from ATCC) (RRID:<u>CVCL_5942</u>) were tested for mycoplasma contamination with the Lookout mycoplasma PCR detection kit (Sigma-Aldrich, St. Louis, MO). Mycoplasma negative HT29-18N2 cells were used for the experiments presented here.

### Patient-derived organoid (PDO)

Samples from patients were kindly provided by MARBiobank and IdiPAZ Biobank, integrated in the Spanish Hospital Biobanks Network (RetBioH; www.redbiobancos.es). Informed consent was obtained from all participants and protocols were approved by institutional ethical committees. For the PDO generation, primary tumour from was disaggregated in 1.5 mg/mL collagenase II (Sigma) and 20 μg/mL hyaluronidase (Sigma) after 40min of incubation at 37ºC, filtered in 100 μm cell strainer and seeded in 50 μL Matrigel in 24-well plates, as previously described (29).

After polymerization, 450 μL of complete medium was added (DMEM/F12 plus penicillin (100 U/mL) and streptomycin (100 μg/mL), 100 μg/mL Primocin, 1X N2 and B27, 10mM Nicotinamide; 1.25 mM N-Acetyl-L-cysteine, 100 ng/mL Noggin and 100 ng/mL R-spondin-1, 10 μM Y-27632, 10 nM PGE2, 3 μM SB202190, 0.5 μM A-8301, 50 ng/mL EGF and 10 nM Gastrin I). Tumour spheres were collected and digested with an adequate amount of trypsin to single cells and re-plated in culture. Cultures were maintained at 37°C, 5% CO_2_ and medium changed every week.

For dose-response curves, 600 single PDO cells were plated in 96-well plates in 10 μL Matrigel with 100 μL of complete medium. After 6 days in culture, the growing PDO was treated with combinations of 5-FU+iri. plus SN6 or 5-FU+iri. After 72 hours of treatment, we measured the cell viability using the CellTiter-Glo 3D Cell Viability Assay following manufacturer’s instructions in an Orion II multiplate luminometer. The mutation identified in the PDO used in this study was KRAS G12D (66.43%).

### Differentiation of HT29-18N2 cells

HT29-18N2 cells were differentiated to goblet cells as described previously (5) Briefly, cells were seeded in complete growth medium (DMEM complemented with 10% FCS and 1% P/S), and the following day (Day zero: D-0), the cells were placed in PFHMII protein free medium (GIBCO, ThermoFisher Scientific, Waltham, MA, USA). After 3 days (D-3), medium was replaced with fresh PFHMII medium and cells grown for 3 additional days. At day 6 (D-6) cells were trypsinized and seeded for the respective experiments in complete growth medium followed by incubation in PFHMII medium at day 7 (D-7). All experimental procedures were performed at day 9 (D-9).

### Survival curves

HT29-18N2 cells (KD/OV) were seeded in 96-well plates with Dulbecco’s modified Eagle’s medium. Confluent cells were treated with combination of 5-Fluorouracil + Irinotecan for 72h at indicated concentrations. Cell viability was determined using the CellTiter-Glo 3D Cell Viability Assay (Promega) following the manufacturer’s instructions in an Orion II multiplate luminometer (Berthold detection systems).

### Western Blot

Cells were lysed 20 min at 4ºC in 500μL of PBS plus 0.5% Triton X-100, 1mM EDTA, 100mM Na-orthovanadate, 0.25mM phenylmethylsulfonyl fluoride, and complete protease inhibitor cocktail (Roche). Lysates were analysed by WB using standard SDS-polyacrylamide gel electrophoresis (SDS-PAGE) techniques. Briefly, protein samples were boiled in Laemmli buffer, run in polyacrylamide gels, and transferred onto polyvinylidene fluoride (PVDF) membranes (Millipore). The membranes were incubated overnight at 4 °C with the appropriate primary antibodies. After being washed, the membranes were incubated with specific secondary horseradish peroxidase-linked antibodies from Dako and visualized using the enhanced chemiluminescence reagent from Amersham.

### Comet assay

HT29-M6 cells were plated in 12-well plates with Dulbecco’s modified Eagle’s medium. 4 days after arriving to 100% confluency, cells were treated with 5-FU+iri. (25 µg/mL 5-Fu + 10 µg/mL Irinotecan) alone or in combination with secretion inhibitor Benzamil 20µM for 72h. Comet assays were performed using CometAssay Trevigen Kit (4250-050-K) following the manufacturer’s instructions. Pictures were taken using a Nikon Eclipse Ni-E epifluorescence microscope and tail moment was calculated using the OPENCOMET plugin for Image J.

### Cell viability assay

HT29-M6 cells were plated in 6-well plates with Dulbecco’s modified Eagle’s medium. After 4 days in postconfluency, CRC cells were treated with 5-FU+iri. (10 µg/mL 5-Fu + 4 µg/mL Irinotecan) alone or in combination with secretion inhibitor Benzamil 20µM. Cell viability was determined by flow cytometry 24h or 72h after treatment, staining cells with DAPI (5μg/mL) and measuring alive cells in the LSR-Fortessa analyzer.

### Microscopy and colocalization analysis

HT29-M6 cells were plated in 6-well plates with Dulbecco’s modified Eagle’s medium. After being 7 days at postconfluency, cells were treated with 5-FU+iri. (50µg/mL 5-Fu + 20µg/mL Irinotecan) alone or in combination with secretor inhibitor SN-6 10µM for 24h. Then cells were directly fixed with 4% paraformaldehyde, permeabilized with 0,3% Triton X-100 (Pierce), washed and incubated overnight with corresponding primary antibodies. The samples were developed with TSA™ Plus Cyanine 3/Fluorescein System (PerkinElmer) and mounted in ProLong™ Diamond Antifade Mountant plus DAPI (Thermo Scientific).

Images were acquired using an inverted Leica SP8 or Leica SP5 confocal microscope with a 63x Plan Apo NA 1.4 objective and analysed using ImageJ ((FIJI, version 2.0.0-rc-43/1.51 g) (21). For detection of the respective fluorescence emission, the following laser lines were applied: DAPI, 405 nm; and Alexa Fluor 555, 561 nm; Alexa Fluor 647, 647 nm. Two-channel colocalization analysis was performed using ImageJ, and the Manders’ correlation coefficient was calculated using the plugin JaCop (30).

### Statistical analysis

All data are means ± SEM. In all cases a D’Agostino– Pearson omnibus normality test was performed before any hypothesis contrast test. Statistical analysis and graphics were performed using GraphPad Prism 6 (RRID:SCR_002798). For data that followed normal distributions, we applied either Student’s t test or one-way analysis of variance (ANOVA) followed by Tukey’s post hoc test. For data that did not fit a normal distribution, we used Mann–Whitney’s unpaired t test and nonparametric ANOVA (Kruskal–Wallis) followed by Dunn’s post hoc test. Criteria for a significant statistical difference were: *p<0.05; **p<0.01.

## ACKNOWLEDGMENTS

We thank all members of the Malhotra and Espinosa Lab for valuable discussions. Fluorescence microscopy was performed at the Advanced Light Microscopy Unit at the CRG, Barcelona. V. Malhotra is an Institució Catalana de Recerca i Estudis Avançats professor at the Centre for Genomic Regulation. This work was funded by grants from the Spanish Ministry of Economy and Competitiveness (BFU2013-44188-P to VM) and FEDER Funds, and Instituto de Salud Carlos III (PI19-00013 to LE). We acknowledge support of the Spanish Ministry of Economy and Competitiveness, through the Programmes “Centro de Excelencia Severo Ochoa 2013-2017” (SEV-2012-0208 & SEV-2013-0347) and Maria de Maeztu Units of Excellence in R&D (MDM-2015-0502). This work reflects only the authors’ views, and the EU Community is not liable for any use that may be made of the information contained therein.

## AUTHORS’ CONTRIBUTION

GCR, MI, LI and VM conceived and designed the research studies. GCR, JAM, TLJ and MG conducted the experiments. GCR, JAM, TLJ, LI and VM analyzed the data. GCR, JAM, TLJ, LI and VM draft the manuscript. All authors read and approved the final manuscript.

## REFERENCES

1. Sheehan, J. K., Kirkham, S., Howard, M., Woodman, P., Kutay, S., Brazeau, C., Buckley, J., and Thornton, D. J. (2004) Identification of molecular intermediates in the assembly pathway of the MUC5AC mucin. J. Biol. Chem. 279, 15698–15705

2. Thornton, D. J., Rousseau, K., and McGuckin, M. A. (2008) Structure and function of the polymeric mucins in airways mucus. Annu. Rev. Physiol. 70, 459–486

3. Adler, K. B., Tuvim, M. J., and Dickey, B. F. (2013) Regulated Mucin Secretion from Airway Epithelial Cells. Front Endocrinol (Lausanne). 10.3389/fendo.2013.00129

4. Rossi, A. H., Sears, P. R., and Davis, C. W. (2004) Ca2+ dependency of “Ca2+-independent” exocytosis in SPOC1 airway goblet cells. J. Physiol. (Lond.). 559, 555–565

5. Mitrovic, S., Nogueira, C., Cantero-Recasens, G., Kiefer, K., Fernández-Fernández, J. M., Popoff, J.-F., Casano, L., Bard, F. A., Gomez, R., Valverde, M. A., and Malhotra, V. (2013) TRPM5-mediated calcium uptake regulates mucin secretion from human colon goblet cells. Elife. 2, e00658

6. Cantero-Recasens, G., Butnaru, C. M., Brouwers, N., Mitrovic, S., Valverde, M. A., and Malhotra, V. (2019) Sodium channel TRPM4 and sodium/calcium exchangers (NCX) cooperate in the control of Ca2+-induced mucin secretion from goblet cells. J. Biol. Chem. 294, 816–826

7. Cantero-Recasens, G., Butnaru, C. M., Valverde, M. A., Naranjo, J. R., Brouwers, N., and Malhotra, V. (2018) KChIP3 coupled to Ca2+ oscillations exerts a tonic brake on baseline mucin release in the colon. Elife. 10.7554/eLife.39729

8. Kesimer, M., Ehre, C., Burns, K. A., Davis, C. W., Sheehan, J. K., and Pickles, R. J. (2013) Molecular organization of the mucins and glycocalyx underlying mucus transport over mucosal surfaces of the airways. Mucosal Immunol. 6, 379–392

9. Pelaseyed, T., Bergström, J. H., Gustafsson, J. K., Ermund, A., Birchenough, G. M. H., Schütte, A., van der Post, S., Svensson, F., Rodríguez-Piñeiro, A. M., Nyström, E. E. L., Wising, C., Johansson, M. E. V., and Hansson, G. C. (2014) The mucus and mucins of the goblet cells and enterocytes provide the first defense line of the gastrointestinal tract and interact with the immune system. Immunol. Rev. 260, 8–20

10. Kufe, D. W. (2009) Mucins in cancer: function, prognosis and therapy. Nat. Rev. Cancer. 9, 874–885

11. Luo, C., Cen, S., Ding, G., and Wu, W. (2019) Mucinous colorectal adenocarcinoma: clinical pathology and treatment options. Cancer Commun (Lond). 10.1186/s40880-019-0361-0

12. Hugen, N., Brown, G., Glynne-Jones, R., de Wilt, J. H. W., and Nagtegaal, I. D. (2016) Advances in the care of patients with mucinous colorectal cancer. Nat Rev Clin Oncol. 13, 361–369

13. Cutsem, E. V., Cervantes, A., Adam, R., Sobrero, A., Krieken, J. H. V., Aderka, D., Aguilar, E. A., Bardelli, A., Benson, A., Bodoky, G., Ciardiello, F., D’Hoore, A., Diaz-Rubio, E., Douillard, J.-Y., Ducreux, M., Falcone, A., Grothey, A., Gruenberger, T., Haustermans, K., Heinemann, V., Hoff, P., Köhne, C.-H., Labianca, R., Laurent-Puig, P., Ma, B., Maughan, T., Muro, K., Normanno, N., Österlund, P., Oyen, W. J. G., Papamichael, D., Pentheroudakis, G., Pfeiffer, P., Price, T. J., Punt, C., Ricke, J., Roth, A., Salazar, R., Scheithauer, W., Schmoll, H. J., Tabernero, J., Taïeb, J., Tejpar, S., Wasan, H., Yoshino, T., Zaanan, A., and Arnold, D. (2016) ESMO consensus guidelines for the management of patients with metastatic colorectal cancer. Annals of Oncology. 27, 1386–1422

14. Mayo, C., Lloreta, J., Real, F. X., and Mayol, X. (2007) In vitro differentiation of HT-29 M6 mucus-secreting colon cancer cells involves a trychostatin A and p27(KIP1)-inducible transcriptional program of gene expression. J Cell Physiol. 212, 42–50

15. Abdullah, L. H., Conway, J. D., Cohn, J. A., and Davis, C. W. (1997) Protein kinase C and Ca2+ activation of mucin secretion in airway goblet cells. Am. J. Physiol. 273, L201–210

16. Guinney, J., Dienstmann, R., Wang, X., de Reyniès, A., Schlicker, A., Soneson, C., Marisa, L., Roepman, P., Nyamundanda, G., Angelino, P., Bot, B. M., Morris, J. S., Simon, I. M., Gerster, S., Fessler, E., De Sousa E Melo, F., Missiaglia, E., Ramay, H., Barras, D., Homicsko, K., Maru, D., Manyam, G. C., Broom, B., Boige, V., Perez-Villamil, B., Laderas, T., Salazar, R., Gray, J. W., Hanahan, D., Tabernero, J., Bernards, R., Friend, S. H., Laurent-Puig, P., Medema, J. P., Sadanandam, A., Wessels, L., Delorenzi, M., Kopetz, S., Vermeulen, L., and Tejpar, S. (2015) The consensus molecular subtypes of colorectal cancer. Nat Med. 21, 1350–1356

17. Jorissen, R. N., Gibbs, P., Christie, M., Prakash, S., Lipton, L., Desai, J., Kerr, D., Aaltonen, L. A., Arango, D., Kruhøffer, M., Orntoft, T. F., Andersen, C. L., Gruidl, M., Kamath, V. P., Eschrich, S., Yeatman, T. J., and Sieber, O. M. (2009) Metastasis-Associated Gene Expression Changes Predict Poor Outcomes in Patients with Dukes Stage B and C Colorectal Cancer. Clin Cancer Res. 15, 7642–7651

18. Watanabe, Y., Koide, Y., and Kimura, J. (2006) Topics on the Na+/Ca2+ exchanger: pharmacological characterization of Na+/Ca2+ exchanger inhibitors. J. Pharmacol. Sci. 102, 7–16

19. Colomer, C., Margalef, P., Villanueva, A., Vert, A., Pecharroman, I., Solé, L., González-Farré, M., Alonso, J., Montagut, C., Martinez-Iniesta, M., Bertran, J., Borràs, E., Iglesias, M., Sabidó, E., Bigas, A., Boulton, S. J., and Espinosa, L. (2019) IKKα Kinase Regulates the DNA Damage Response and Drives Chemo-resistance in Cancer. Molecular Cell. 75, 669-682.e5

20. Niu, C.-F., Watanabe, Y., Ono, K., Iwamoto, T., Yamashita, K., Satoh, H., Urushida, T., Hayashi, H., and Kimura, J. (2007) Characterization of SN-6, a novel Na+/Ca2+ exchange inhibitor in guinea pig cardiac ventricular myocytes. Eur. J. Pharmacol. 573, 161–169

21. Schindelin, J., Arganda-Carreras, I., Frise, E., Kaynig, V., Longair, M., Pietzsch, T., Preibisch, S., Rueden, C., Saalfeld, S., Schmid, B., Tinevez, J.-Y., White, D. J., Hartenstein, V., Eliceiri, K., Tomancak, P., and Cardona, A. (2012) Fiji: an open-source platform for biological-image analysis. Nat. Methods. 9, 676–682

22. Podhorecka, M., Skladanowski, A., and Bozko, P. (2010) H2AX Phosphorylation: Its Role in DNA Damage Response and Cancer Therapy. J Nucleic Acids. 2010, 920161

23. Cone, R. A. (2009) Barrier properties of mucus. Adv. Drug Deliv. Rev. 61, 75–85

24. Zhu, Y., Abdullah, L. H., Doyle, S. P., Nguyen, K., Ribeiro, C. M. P., Vasquez, P. A., Forest, M. G., Lethem, M. I., Dickey, B. F., and Davis, C. W. (2015) Baseline Goblet Cell Mucin Secretion in the Airways Exceeds Stimulated Secretion over Extended Time Periods, and Is Sensitive to Shear Stress and Intracellular Mucin Stores. PLoS ONE. 10, e0127267

25. Pothuraju, R., Rachagani, S., Krishn, S. R., Chaudhary, S., Nimmakayala, R. K., Siddiqui, J. A., Ganguly, K., Lakshmanan, I., Cox, J. L., Mallya, K., Kaur, S., and Batra, S. K. (2020) Molecular implications of MUC5AC-CD44 axis in colorectal cancer progression and chemoresistance. Mol Cancer. 19, 37

26. Jin, W., Liao, X., Lv, Y., Pang, Z., Wang, Y., Li, Q., Liao, Y., Ye, Q., Chen, G., Zhao, K., and Huang, L. (2017) MUC1 induces acquired chemoresistance by upregulating ABCB1 in EGFR-dependent manner. Cell Death Dis. 8, e2980

27. Kamnerdsupaphon, P., Lorvidhaya, V., Chitapanarux, I., Tonusin, A., and Sukthomya, V. (2007) FOLFIRI chemotherapy for metastatic colorectal cancer patients. J Med Assoc Thai. 90, 2121–2127

28. Iwamoto, T., Inoue, Y., Ito, K., Sakaue, T., Kita, S., and Katsuragi, T. (2004) The exchanger inhibitory peptide region-dependent inhibition of Na+/Ca2+ exchange by SN-6 [2-[4-(4-nitrobenzyloxy)benzyl]thiazolidine-4-carboxylic acid ethyl ester], a novel benzyloxyphenyl derivative. Mol. Pharmacol. 66, 45–55

29. Sato, T., Stange, D. E., Ferrante, M., Vries, R. G. J., Van Es, J. H., Van den Brink, S., Van Houdt, W. J., Pronk, A., Van Gorp, J., Siersema, P. D., and Clevers, H. (2011) Long-term expansion of epithelial organoids from human colon, adenoma, adenocarcinoma, and Barrett’s epithelium. Gastroenterology. 141, 1762–1772

30. Bolte, S., and Cordelières, F. P. (2006) A guided tour into subcellular colocalization analysis in light microscopy. J Microsc. 224, 213–232

